# Divergent Volatile Metabolomes and Flavor Attributes in Rice Fermented by *Aspergillus oryzae* and *Aspergillus flavus*

**DOI:** 10.1101/2025.07.07.663597

**Authors:** Colin M. McCarthy, Dasol Choi, Alissa A. Nolden, Eric A. Decker, Jae-Hyuk Yu, John G. Gibbons

## Abstract

The generally recognized as safe (GRAS) fungus *Aspergillus oryzae* has been used for millennia in the production of traditional Asian fermented foods and beverages. This prolonged domestication has led to phenotypic adaptations that distinguish *A. oryzae* from its wild relative*, Aspergillus flavus.* While genomic and phenotypic differences between these species have been partially characterized, their comparative production of volatile compounds during food fermentation remains poorly understood. Here, we evaluated alpha-amylase activity, aflatoxin production, taste attributes (via electronic tongue), and volatile profiles (via dynamic headspace gas chromatography–mass spectrometry) of the food-grade strain *A. oryzae* RIB40 and two wild *A. flavus* strains, NPK13tox and AflaGuard*. A. oryzae* RIB40 exhibited significantly higher alpha-amylase activity during rice fermentation, and only *A. flavus NPK13tox* produced aflatoxin. Sensory analysis revealed that rice fermented by *A. oryzae* RIB40 exhibited more favorable taste attributes, including significantly lower astringency, aftertaste, and bitterness, as well as significantly higher richness (defined as umami aftertaste). Additionally*, A. oryzae* RIB40 produced a greater number and higher concentrations of volatile compounds, with 20 compounds significantly elevated compared to rice fermented by *A. flavus.* Many of these volatiles, including 2-methyl-3-buten-2-ol, 3-octen-2-one, 2-methyl-butanal, and 3-methyl-butanal, are associated with pleasant sensory attributes and have previously been linked to *A. oryzae-*fermented food*s.* These findings suggest that the volatilome of *A. oryzae* RIB40 has been shaped by domestication to produce a more desirable sensory profile, enriched in alcohols, aldehydes, ketones, and heterocyclic compounds contributing fruity, umami, and malty notes.

## Introduction

Traditional fermented foods and beverages are a staple of many Asian diets. For instance, nearly 20% of the total weight and energy of Japanese dietary intake is from fermented foods [1]. *Aspergillus oryzae* is a filamentous mold and one of the most commonly used microbes for fermented food production in Asia (*e.g.* miso, soy sauce, sake and amazake). In soy-based fermentations, *A. oryzae* is used primarily for its high proteolytic activity, while in rice-based fermentations *A. oryzae* is used for both proteolytic and amylolytic activity [2, 3]. *A. oryzae* was domesticated over thousands of years of use in food fermentation from its wild progenitor *Aspergillus flavus* [4, 5], which is a major agricultural pest and the main producer of aflatoxin contamination in stored seeds and grains [6]. As a result of domestication, *A. oryzae* lost its ability to produce some toxic secondary metabolites (including aflatoxin, cyclopiazonic acid, and aflatrem [2, 4, 7, 8]) and adapted to the high starch environment of rice by increasing alpha-amylase production through gene duplications of the alpha-amylase encoding gene [9, 10]. While the genetic differences between *A. oryzae* and *A. flavus* are well documented, far less is understood about how domestication shaped the sensory characteristics of *A. oryzae*.

During fermentation, *A. oryzae* produces a variety of extracellular enzymes to digest proteins, carbohydrates, and lipids into sugars, amino acids and free fatty acids that are subsequently reabsorbed and used as energy. These metabolites also significantly influence the volatile profile and sensory characteristics of the fermented food product. For instance, *A. oryzae* produces lipases during doenjang (soy paste) fermentation, and lipase activity shows strong associations with the formation of longIllchain fatty acid esters that are essential for the final aroma and flavor [11]. Additionally, in soy sauce fermentation, *A. oryzae* increases certain volatile compounds that impart “musty” and “soy-sauce-like” odors [12]. Importantly, Zhao et al. (2015) showed substantial variation in volatile compound profiles (e.g. levels of ketones, aldehydes, alcohols and esters) between two *A. oryzae* strains during soy fermentation, which indicates that genetic differences within *A. oryzae* can directly result in distinct volatile profiles [13].

Less is understood regarding the volatile, sensory, and enzymatic differences of foods fermented by these two strains, *A. oryzae* and *A. flavus*. Addressing these differences could provide insight into metabolic shifts impacted by *A. oryzae*’s long-term adaptation to food substrates. Rank *et al.* (2012) performed perhaps the only direct comparative metabolic study of the *A. oryzae* and *A. flavus* reference strains (RIB40 and NRRL 3357, respectively) using an LC-MS/MS, and found species-specific chemical profiles, including the absence of secondary metabolite end-products in *A. oryzae* [7]. This result suggests *A. oryzae* and *A. flavus* have distinct metabolomic profiles, which likely impact their volatile profiles and sensory characteristics.

Here, we measured aflatoxin production, alpha-amylase activity, volatile profiles (via Dynamic Headspace Gas Chromatography Mass Spectrometry (DH-GC-MS)), and taste attributes (via E-tongue) of the fermentation starter culture *A. oryzae* RIB40 (AO_RIB40), the aflatoxin producer *A. flavus* NPK13tox (AF_NPK13tox), and the non-aflatoxin producer and biocontrol strain *A. flavus* AflaGuard (AF_AflaGuard). The goal of this study was to understand the differences in volatile and taste characteristics between the domesticated food grade starter culture *A. oryzae* and its wild progenitor *A. flavus*.

## Materials and Methods

### *Aspergillus flavus* 13tox genome sequencing and relationship of strains analyzed

We sequenced the genome of AF_NPK13tox to understand its relationship to AO*_*RIB40 and AF_AflaGuard. *A. flavus* 13tox was grown on potato dextrose agar (PDA) at 30°C for 48 hours. DNA was extracted directly from the spores using the method outlined by [14]. DNA concentrations were measured for each extraction using a Qubit fluorometer. PCR-free 150-bp paired-end libraries were prepared and sequenced by Novogene (https://en.novogene.com/) on an Illumina NovaSeq 6000 platform. The raw whole-genome sequencing data for *A. flavus* 13tox can be accessed through the BioProject accession number PRJNA1277813. Raw paired-end fastq files were adapter and quality trimmed using trim_galore [15] with the following parameters: “-q 30”, “--stringency 1”, and “--length 80”. Adapter and quality trimmed reads were assembled using SPAdes [16] using the “--careful” option and K-mer sizes of 55, 75, 95, 105, and 115.

Eleven genome assemblies spanning the diversity of *A. oryzae* and *A. flavus* populations were obtained from NCBI and analyzed with the AF_NPK13tox assembly. These assemblies were *A. flavus* AflaGuard (GCA_012896875.1), OK-A-8-1-S (GCA_024676875.1), NRRL 3357 (GCF_009017415), *A. oryzae* TK14 (GCA_009685055.1), TK31 (GCA_009686725.1), TK49 (GCA_009687085.1), TK56 (GCA_009687225.1), TK57 (GCA_009687245.1), TK60 (GCA_009687305.1), RIB40 (GCF_000184455.2), 14160 (GCA_019097755.1) [2, 5, 9, 17, 18], and *Aspergillus parasiticus* CBS-117618 (GCA_009176385.1). For this analysis, gene models for all 11 genomes were predicted using Augustus v3.4.0 with the following parameters: “--species=aspergillus_oryzae,” “--strand=both,” “--gff3=on,” “--uniqueGeneId=true,” and “--protein=on” [19]. We used Augustus for gene prediction even for genomes with existing annotations to avoid bias from differences in gene model predictions. Single-copy orthologs across the 13 proteomes were identified using OrthoFinder v2.5.5 [20, 21]. The identified single copy ortholog proteins were concatenated and aligned using MAFFT v7.481 [22]. A phylogenetic tree was then constructed from the concatenated protein alignment with IQ-TREE v2.1.3 using 1,000 bootstrap replicates (-B 1000) and the LG amino acid replacement model [23]

### Fungal Isolates

AO*_*RIB40, AF_NPK13tox, and AF_AflaGuard were selected for rice fermentation and subsequent analysis because these strains represent distinct populations within the *A. oryzae*/*A. flavus* species complex (**Figure 1A**).

**Figure 1.**
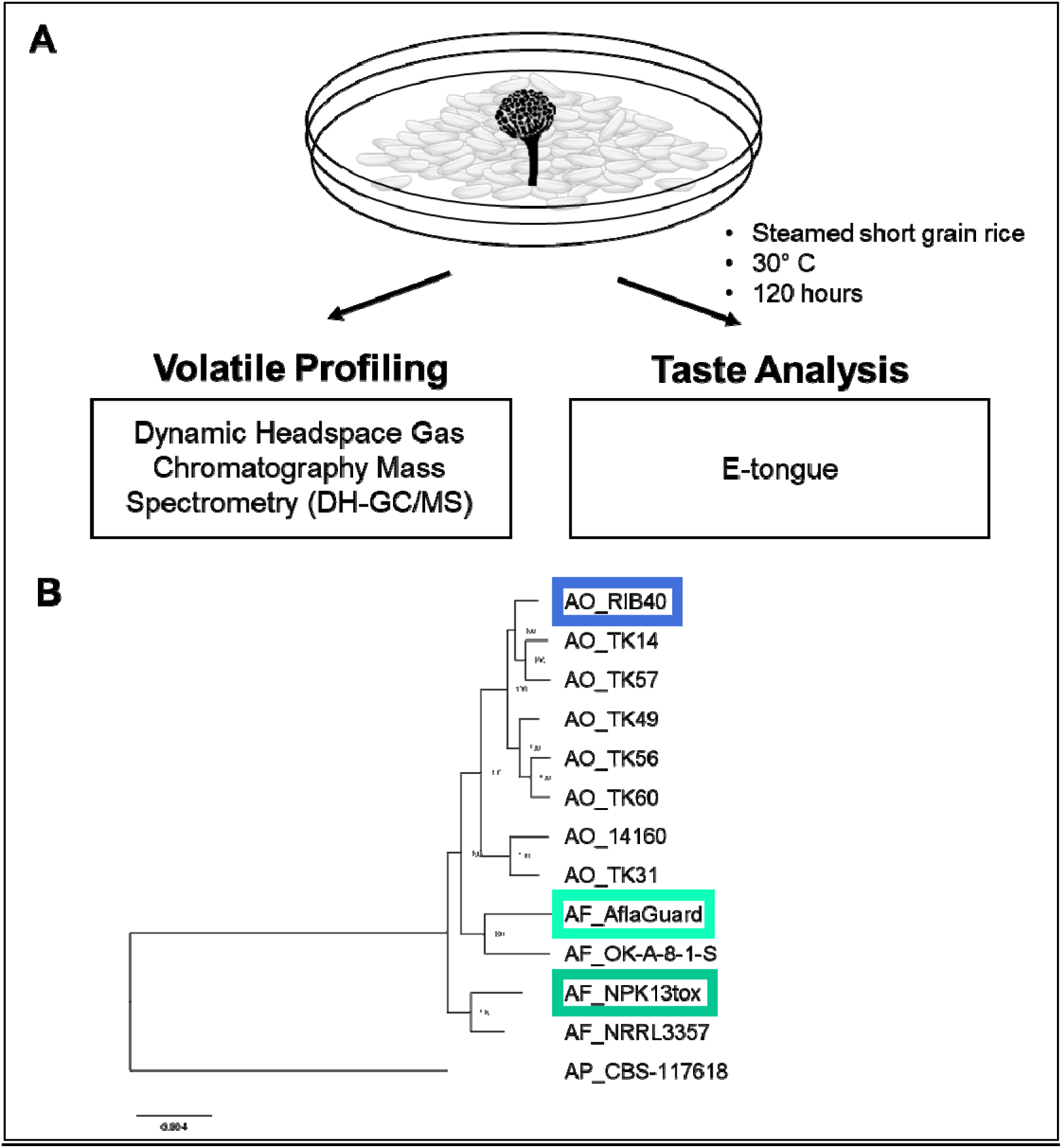
Rice fermentation experimental design and relationship of starter cultures. (A) Schematic of the experimental design, in which starter cultures fermented steamed rice after which the fermented products were subjected to volatile compound profiling via Dynamic Headspace Gas Chromatography Mass Spectrometry (DH-GC/MS), and flavor analysis via electronic tongue (e-tongue). (B) Maximum likelihood phylogenetic tree generated from a concatenated alignment of 8,092 single copy orthologs across 8 *A. oryzae* genomes (AO) and 4 *A. flavus* genomes (AF) and one *A. parasiticus* genome (AP), which was used to root the tree. Node labels represent bootstrap values. Genomes outlined in colored boxes note the strains used for rice fermentation in this study.

### Alpha-amylase activity

We measured alpha amylase activity during rice fermentation across the three fermentation starters, because alpha amylase activity is essential for rice fermentation and is one of the defining features of *A. oryzae* domestication [4, 10]. To measure alpha amylase activity, 12.25 g of fermented rice samples were combined with 10 mL of molecular water, vortexed for 1 minute, and a 2 mL aliquot was withdrawn with analysis. The 2 mL aliquot was fractionated by centrifugation at 1,000 g for 10 min. A 1 mL aliquot of the resulting supernatant was withdrawn and used for alpha amylase quantification. Alpha amylase activity of each strain was evaluated with Ceralpha α-Amylase Assay Kit (cat. No. K-CERA) in biological triplicate. This method utilizes non-reducing end-blocked p-nitrophenyl maltoheptaoside in the presence of an excess of thermostable α-glucosidase. The supernatant was diluted 1:24 with the assay buffer before proceeding with the assay protocol. Absorbance values (405 nm) were recorded with a SpectraMax i3 microplate reader (Molecular Devices, USA). The negative control for this assay was developed by combining all reagents in the same proportions as test samples, but without incubation such that the active enzyme was denatured and thus no longer capable of hydrolyzing starch. This was to confirm that any changes in absorbance (405 nm) were due to enzymatic activity rather than non-enzymatic activity or baseline fluctuations.

### Aflatoxin quantification

Aflatoxin quantification was carried out using a modified protocol adapted from Alshannaq et al. (2018) and Acevedo et al. (2025) [24, 25]. Isolates were grown in slant cultures prepared with 2 mL of potato dextrose (PD) medium in 10 mL glass test tubes. Approximately 10_J conidia were inoculated into each tube using a sterile loop. The tubes were placed at a 45° angle on a rack and incubated at 30° C for 7 days to allow for mycelial growth and toxin production. After incubation, aflatoxins were extracted using an organic solvent method. One mL of chloroform was added directly to each culture tube, thoroughly mixed, and then centrifuged at 5,000 × g to facilitate phase separation. From the chloroform phase, 0.5 mL was carefully collected and left to evaporate at room temperature. The resulting residue was reconstituted in 0.5 mL of High-performance liquid chromatography (HPLC) mobile phase consisting of water, methanol, and acetonitrile (50:40:10, v/v/v). Samples were filtered through a 0.45 μm membrane filter prior to HPLC analysis.

HPLC was used to quantify aflatoxins AFB1, AFB2, AFG1, and AFG2. Analyses were performed on an Agilent 1100 series system equipped with a degasser, autosampler, quaternary pump, and a 1260 Infinity diode array detector (Agilent Technologies, CA, USA). Separation was achieved using a Zorbax Eclipse XDB-C18 column (4.6 mm × 150 mm, 3.5 μm particle size) with a flow rate of 0.8 mL/min. Aflatoxins were detected at a wavelength of 365 nm.

### Culture conditions for e-tongue and DH-GC/MS analysis

Freezer stocks of conidia for each strain were rapidly thawed and cultured on steamed rice 7 days at 30°C in petri dishes, at which time, the plates were flooded with 50% water + 49.9% glycerol + 0.1% tween to harvest conidia. Conidia were then quantified on a haemocytometer and normalized to 1×10^6^ conidia per mL. Rice was prepared for fermentation by autoclaving short grain white rice at 170% hydration for 15 minutes at 121°C. 12 g of sterilized short grain rice was inoculated with 1 mL of normalized conidia stocks (∼1 million conidia) in 60 mm petri dishes sealed with parafilm and incubated for 48 hours at 30°C. Fermentations were performed in duplicate. As a negative control we also performed the same experiment but added 1 mL of the conidia harvesting solution (50% water + 49.9% glycerol + 0.1% tween) without conidia (rice only). At the end of the 48 hours period, samples were transferred to 50 mL conical tubes, 25 mL of water was added, and samples were vortexed for 60 s. This fermented rice/water slurry closely resembles amazake and early stage sake fermentation and was thus considered a suitable testbed for organoleptic quality analysis. Samples were flash frozen in liquid nitrogen and shipped to Medallion Labs (Minneapolis, MN) for E-tongue and DHS-GC-MS analysis using an unfractionated aliquot of the fermented rice/water slurry. Our overall experimental design is depicted in **Figure 1A**.

### E-tongue

Traditional descriptive sensory testing with human subjects was not possible because AF_NPK13tox is an aflatoxin producer. Thus, we analyzed our samples via e-tongue, which measures flavor attributes by using an array of chemical sensors combined with pattern recognition algorithms to detect and differentiate flavor-related compounds in a liquid sample. E-tongue analysis assessed these compounds and provided an interpretation of how these might impact eight different flavor profiles (i.e., umami, aftertaste richness of umami, saltiness, sourness, bitterness, aftertaste of bitterness, astringency, and aftertaste of astringency). These attributes are based on a relative comparison to internally developed references. Data were reported as relative comparisons to the control, using a scale where 1 unit represents a level of differential sensitivity equivalent to an estimated 20% difference in taste. Two biological replicates were analyzed for each fermentation, and one sample was analyzed for the unfermented rice control. For each flavor attribute, we conducted an ANOVA between the fermentation starter strains assuming a p-value < 0.01 cutoff. For comparisons in which p-values < 0.01, we conducted post hoc Tukey Kramer HSD tests to compare pairwise differences between samples, imposing a p-value < 0.01 cutoff.

### DHS-GC-MS

Samples were analyzed as received by dynamic headspace on an Agilent 5977 MSD coupled to an Agilent 7890A gas chromatography unit with a Gerstel multipurpose sampler (Agilent, USA). Compounds were identified and matched by a spectrum library. Quantitation of compounds present was determined by comparison of their response to that of internal standard added to each sample. Prior to testing, samples were quantified as compound profiles in Parts Per Million (ppm). All sample processing, compound identification and quantification were performed at Medallion Labs (Minneapolis, MN). For each compound, we conducted an ANOVA between the fermentation starters strains assuming a p-value < 0.01 threshold. For comparisons in which p-values < 0.01, we conducted post hoc Tukey Kramer HSD tests to compare pairwise differences between samples, imposing a p-value < 0.01 cutoff.

## Results

### Fermentation strains are from phylogenetically distinct groups

We analyzed the phylogenetic relationship of the three rice fermentation starters (*A*O_RIB40, AF_AflaGuard and AF_NPK13tox) in relation to 7 additional *A. oryzae* genomes, 2 additional *A. flavus* genomes that are representative of the major populations based on previously published work [4, 5, 9, 18, 26], and a strain of *A. parasiticus* which was used as an outgroup. The genome of AF_NPK13tox was sequenced and assembled for the current study (BioProject ID PRJNA1277813), while all other genome assemblies were publicly available (see methods for accession numbers). We used OrthoFinder to identify single copy orthologs [20, 21], and then built a maximum likelihood phylogenetic tree based on a protein alignment of 8,092 single copy orthologs (which included 4,445,407 amino-acid sites and 48,990 parsimony informative sites). The phylogenetic tree showed three phylogenetically distinct clades of *A. oryzae*, and two clades of *A. flavus* (**Figure 1A**), consistent with previous studies. Importantly, the three fermentation strains resided in distinct clades.

### Alpha-amylase activity is significantly higher in *A. oryzae* RIB40

We measured alpha-amylase activity at the end of our fermentation because variability in alpha-amylase production could impact volatile compound formation (*e.g.* higher sugar concentrations could lead to more ethanol production by yeasts which in turn could increase the pool of substrates for ester synthesis), and taste perception (*e.g.* higher alpha-amylase activity would increase sugar content and sweetness of the fermentation product) [27]. Unsurprisingly, alpha-amylase activity was significantly different between isolates (ANOVA, p-value = 5.2e-6) and significantly higher in AO_RIB40 compared to the *A. flavus* isolates (Tukey-Kramer: AO_RIB40 vs. AF_NPK13tox: p-value = 1.0e-4 and AO_RIB40 vs. AF_AflaGuard: p-value = 8.6e-6).

### Only *A. flavus* NPK13tox produces aflatoxins

We quantified the production of aflatoxins (AFB₁, AFB₂, AFG₁, and AFG₂) by AO_RIB40, AF_NPK13tox, and AF_AflaGuard under culture conditions known to promote aflatoxin biosynthesis [24, 25]. Aflatoxins were not detected in cultures of AO_RIB40 or AF_AflaGuard. In contrast, AF_NPK13tox produced approximately 97 ppm AFB₁ and 34 ppm AFG₁. These results are consistent with prior knowledge: AF_NPK13tox is closely related to the aflatoxigenic strain *A. flavus* NRRL 3357 (**Figure 1A**), whereas AO_RIB40 and AF_AflaGuard harbor well-documented loss-of-function mutations in the aflatoxin biosynthetic gene cluster [4, 25].

### E-tongue analysis for *A. oryzae* RIB40 fermentation has more favorable profile

Fermented rice water slurries underwent e-tongue analysis to measure and compare flavor attributes between starter cultures. Compounds known to associate with perceived saltiness, sourness, umami, umami aftertaste, astringency, aftertaste-A (astringency aftertaste), bitterness and aftertaste-B (bitter aftertaste) were measured for the unfermented rice control, and the biological replicates of each fermentation starter culture. Four of the 8 comparisons were statistically significant (ANOVA, all p-values ≤ 0.002, **Figure 3**). Astringency is a tactile sensation perceived as dryness, roughness, and a tightening feeling in the mouth, often accompanied by a slight contraction of the tongue and oral tissues, and it is often accompanied with acidity or bitterness sensations [28]. Interestingly, astringency and aftertaste-A was significantly lower in the AO_RIB40 fermentation than in the *A. flavus* fermentations (**Figure 3**). Additionally, bitterness was substantially lower in all samples relative to the rice control and significantly lower in the AO_RIB40 fermentation compared to the *A. flavus* fermentations which were themselves significantly different (**Figure 3**). Lastly, umami aftertaste, described as the richness, complexity, and full bodied, was significantly higher in the AO_RIB40 fermentation compared to the *A. flavus* fermentations which, were also significantly different from each other (**Figure 3**).

**Figure 2.**
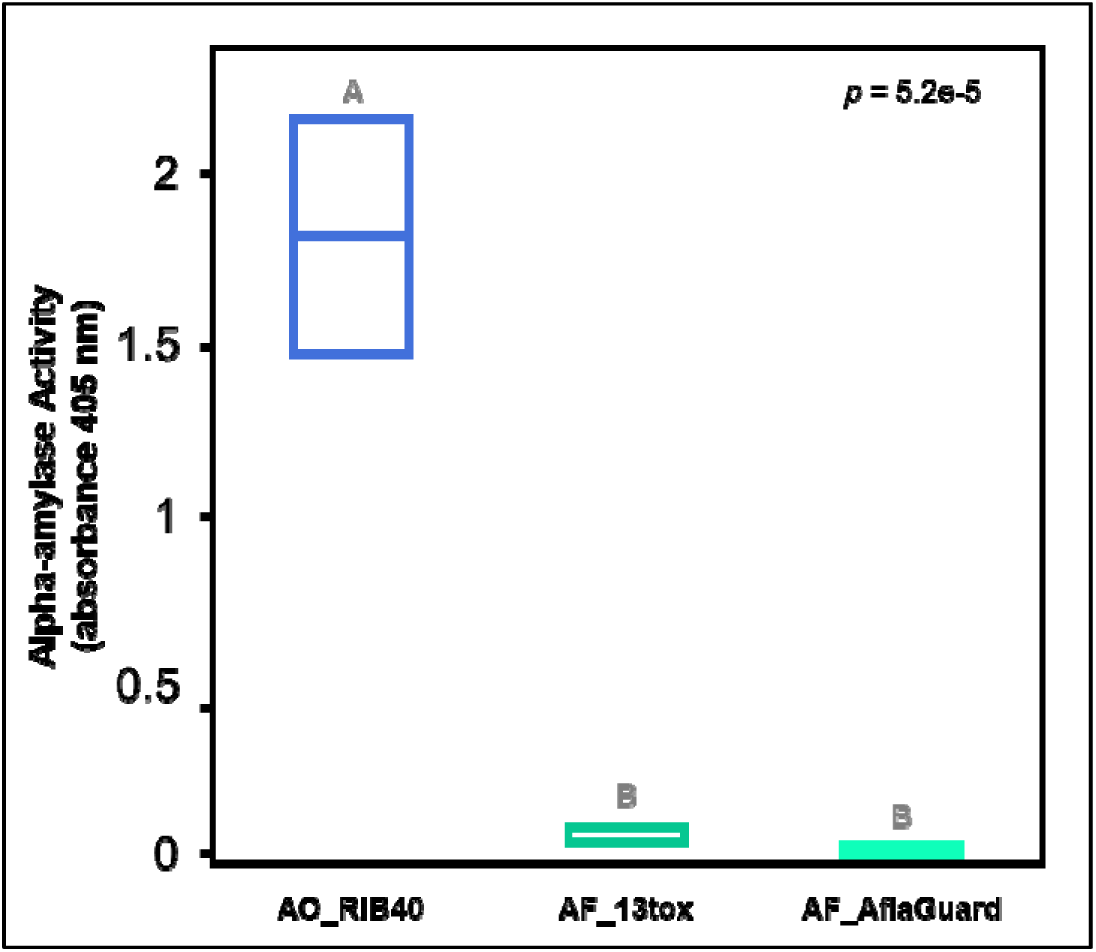
Alpha-amylase activity during rice fermentation. Alpha-amylase activity was quantified in the rice fermentation with the Ceralpha α-Amylase Assay Kit (cat. No. K-CERA) in biological triplicate. Box plots for each strain (x-axis) represent alpha-amylase activity (y-axis). The ANOVA p-value is reported, and letters above box plots signify statistically different groups based on post hoc Tukey Kramer HSD tests. AO_RIB40 has higher alpha-amylase activity compared to both *A. flavus* strains during rice fermentation.

**Figure 3.**
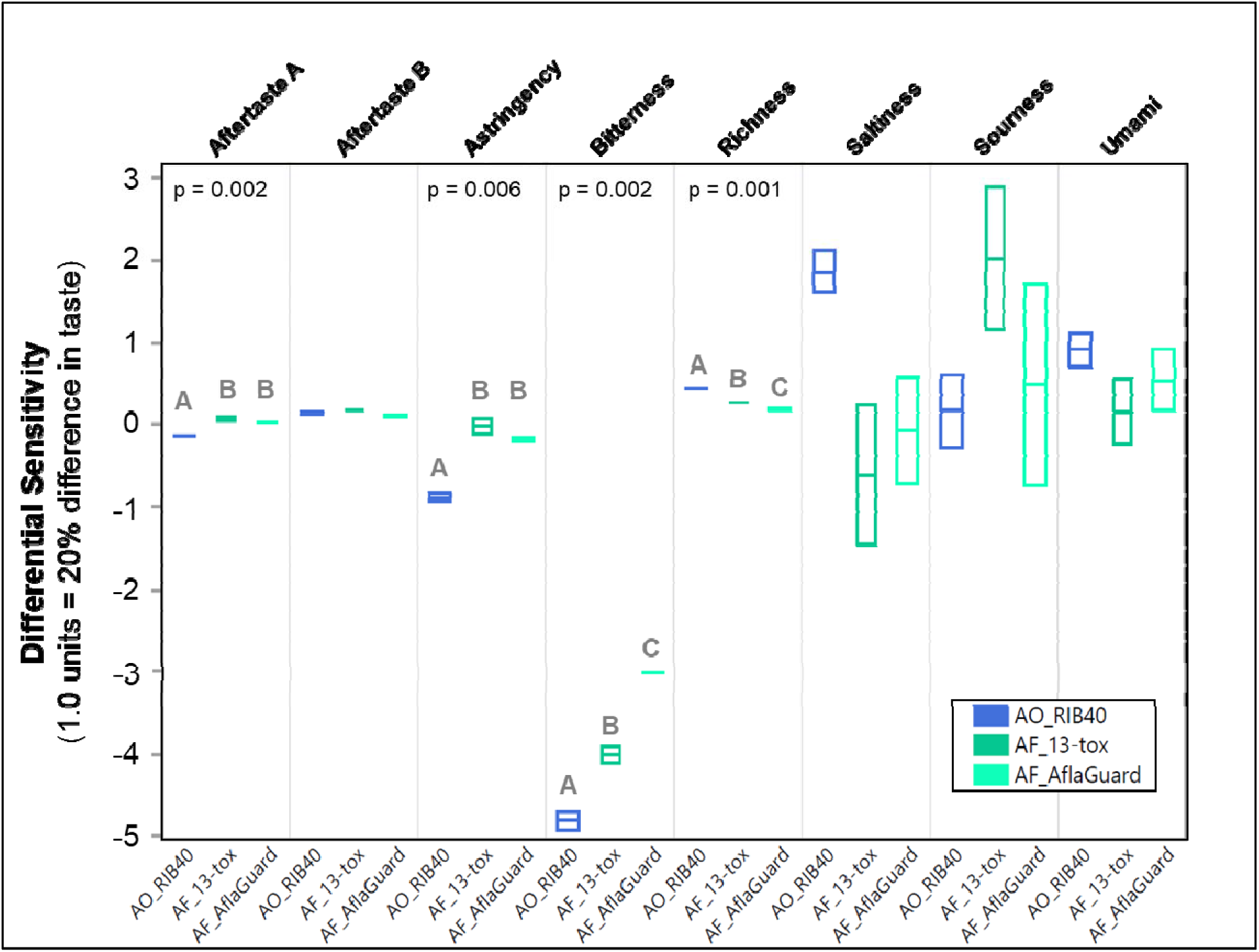
E-tongue analysis of rice fermentation using different starter cultures. Box plots of 8 taste attributes measured by e-tongue (y-axis) across rice fermented by AO_RIB40, AF_NPK13tox, and AF_AflaGuard (x-axis). Box plots represent measurements from three biological replicates per starter culture. The ANOVA p-value is reported for taste attributes that were statistically significant via ANOVA across the three starter cultures, and letters above box plots signify statistically different groups based on post hoc Tukey Kramer HSD tests. The Y-axis represents differential sensitivity in which 1 unit represents a 20% difference in taste.

### Volatilome profiles vary significantly between AO and AF samples

We performed DHS-GC-MS of the fermented rice water slurries to identify unique compounds and compare the relative abundances of compounds between starter cultures. We detected 34, 35, 48, and 61 unique compounds in the rice slurry without a starter culture, the AF_NPK13tox fermentation, the AF_AflaGuard fermentation, and the AO_RIB40 fermentation, respectively (**Figure 4A**, **Table 1**). Between the starter culture samples, 30 compounds were detected in all samples, while 1, 4 and 19 compounds were uniquely detected in the AF_NPK13tox, AF_AflaGuard, and the AO_RIB40 fermentations, respectively (**Figure 4B**).

**Figure 4.**
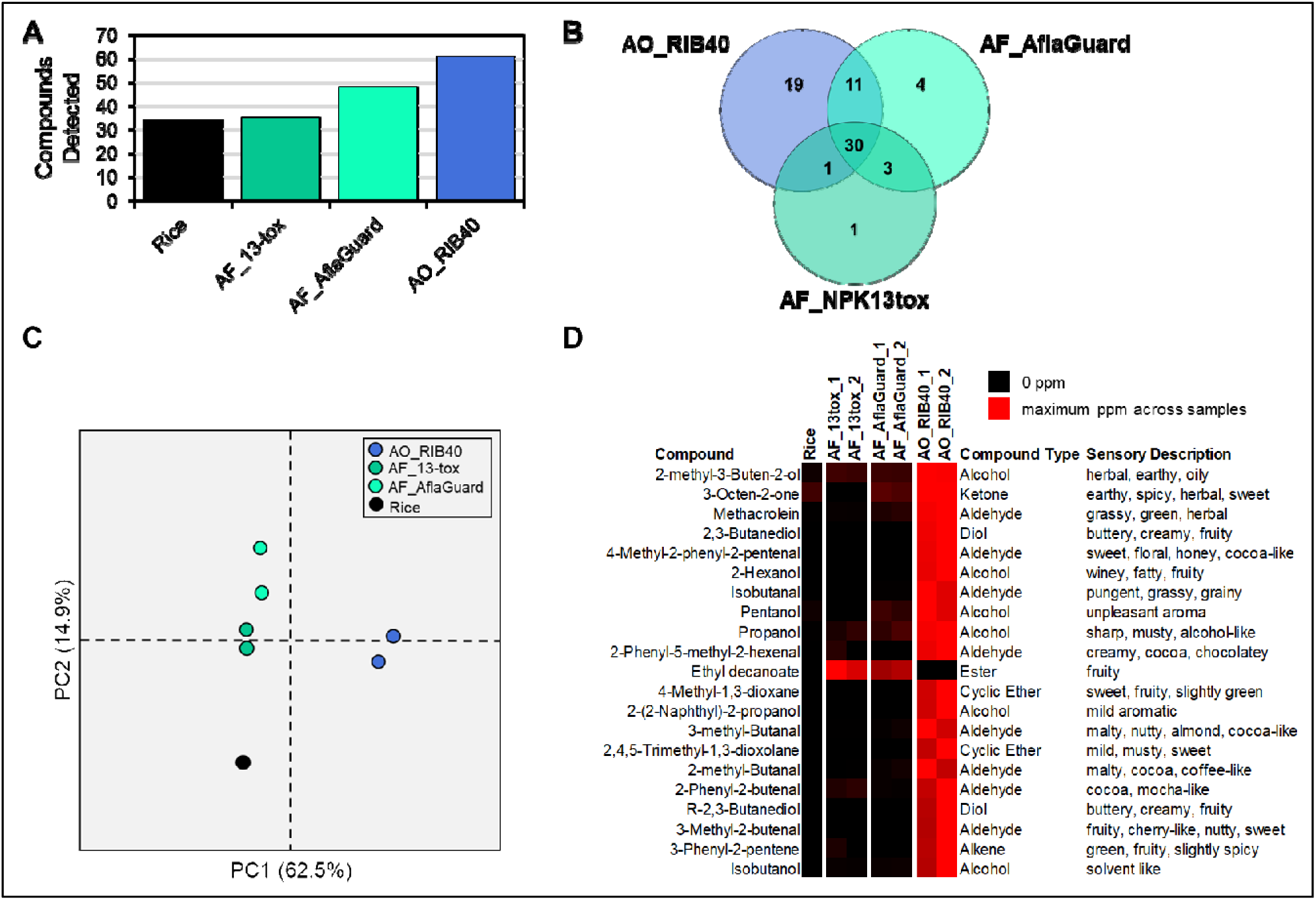
Volatile compound profiling of rice fermentation using different starter cultures. Volatile compound profiles were identified relatively quantified for rice fermentations across biological replicates of AO_RIB40, AF_NPK13tox, and AF_AflaGuard and unfermented rice. (A) Number of unique volatile compounds detected the unfermented and fermented rice samples. (B) Venn diagram of volatile compounds between AO_RIB40, AF_NPK13tox, and AF_AflaGuard rice fermentations. (C) Principal Components Analysis of volatile compound profiles between samples. PC1 separates AO_RIB40 from the other samples and explains 62.5% of variance, while PC2 separates the *A. flavus* samples and rice sample and explains 14.9% of variance. (D) Heatmap depicting the significant differences in volatile compounds abundance between AO_RIB40, AF_NPK13tox, and AF_AflaGuard rice fermentations. The heatmap is relative for each compound and black represents absence of the compound while red represents the maximum PPM value.

**Table 1.**
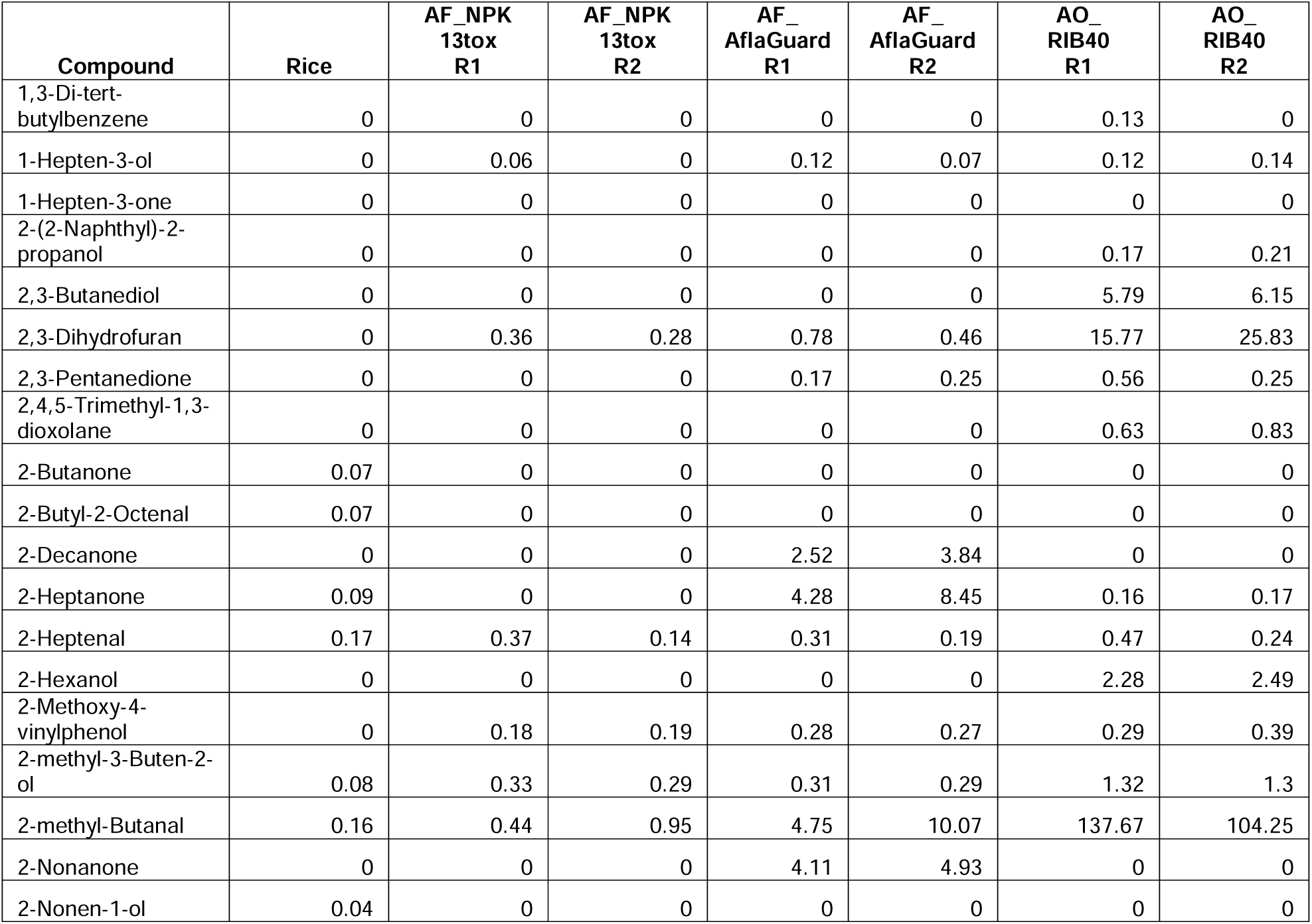
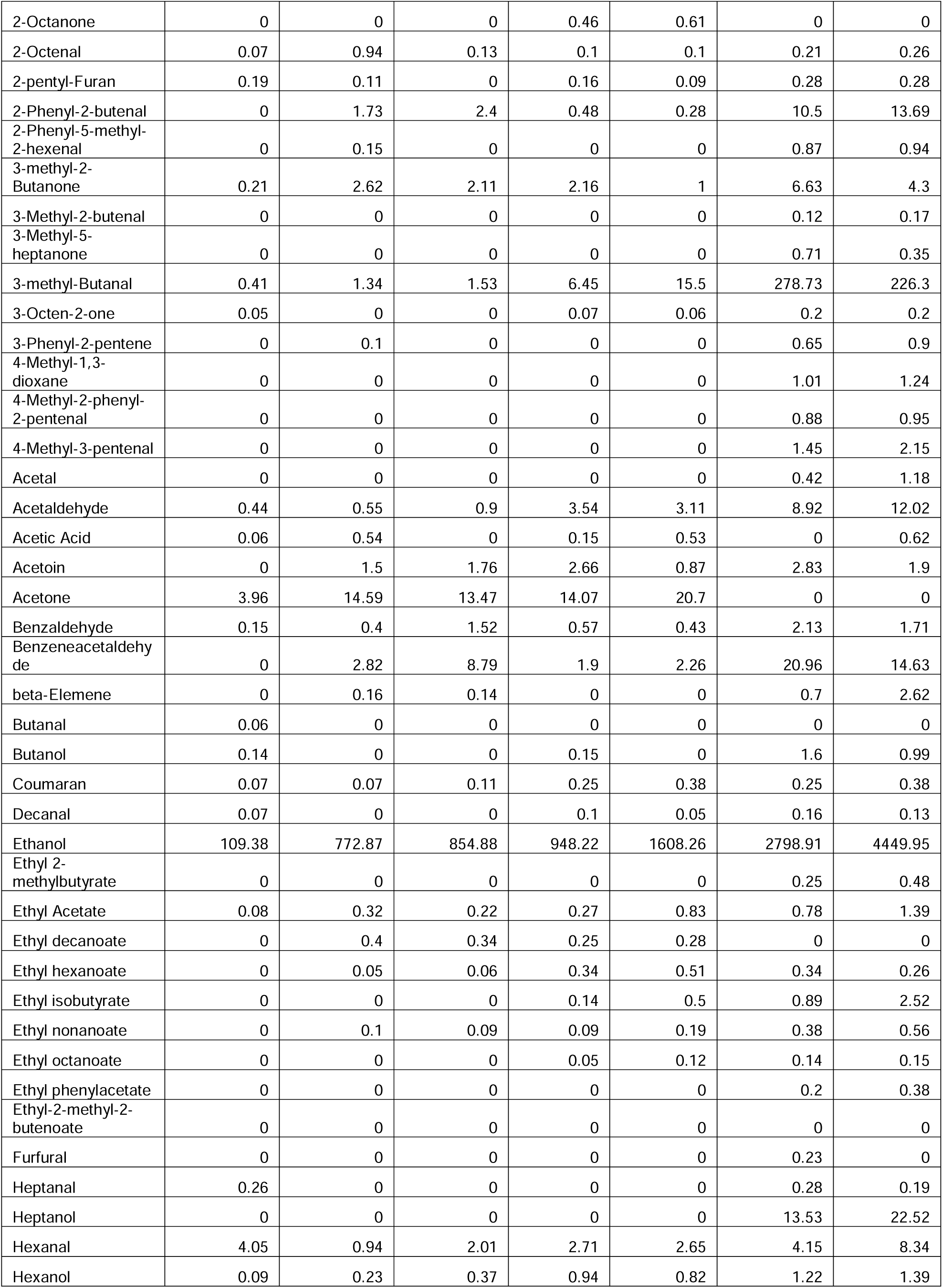
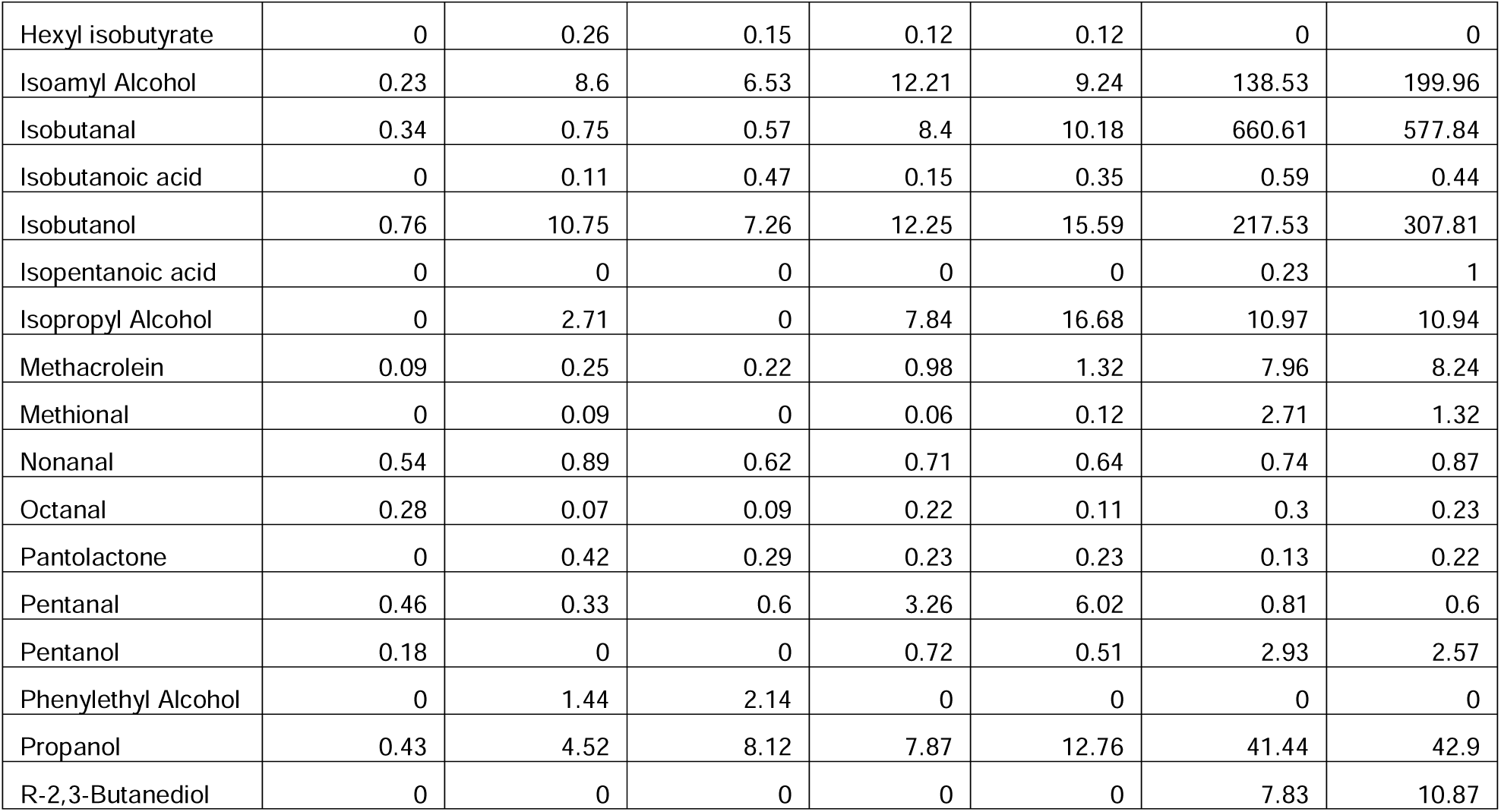
Volatile compounds quantification (parts per million) across unfermented rice, and rice fermented by AO_RIB40, AF_NPK13tox, and AF_AflaGuard. *R1 and R2 represent biological replicates.

To understand how samples volatile profiles were related, we performed a PCA on each sample and replicate using the compound abundance (in parts per million) matrix as input. PC1 explained 62.5% of variance, and separated rice, AF_NPK13tox, and AF_AflaGuard, by AO_RIB40, while PC2 explained 14.9% of variance and separated the AF strains and the rice sample (**Figure 4C**). To test for differences in compound abundance between the three starter cultures, we performed an ANOVA for each compound, imposing a p-value cutoff of 0.01. For compounds where abundance significantly differed between starter cultures, we performed post-hoc Tukey’s HSD tests to identify pairwise differences imposing a cutoff of p-value < 0.01. In total, we identified significant differences between samples for 24 compounds (**Figure 4D**). Only 1 of the 24 compounds, ethyl decanoate, was found at significantly lower abundance in AO_RIB40 in comparison with one or both of the AF starter cultures. Ethyl decanoate, which has been linked to both sweet and vinegar aromas [29], was not detected in the AO_RIB40 fermentations.

The remaining 23 compounds were found at significantly higher abundance in AO_RIB40 than the AF_NPK13tox and AF_AflaGuard (**Figure 4D**). These compounds include 6 alcohols, 8 aldehydes, 1 alkene, 2 cyclic ethers, 2 diols, 2 esters, and 1 ketone. Interestingly, among these 23 compounds, many have sensory descriptions as “*fruity*” (2,3-butanediol, 2-hexanol, 4-methyl-1,3-dioxane, R-2,3-butanediol, 3-methyl-2-butenal and 3-phenyl-2-pentene), “*sweet*” (3-octen-2-one, 4-methyl-2-phenyl-2-pentenal, 4-methyl-1,3-dioxane, 2,4,5-trimethyl-1,3-dioxolane and 3-methyl-2-butenal) and either “*cocoa*” or “*cocoa-like*” (4-methyl-2-phenyl-2-pentenal, 2-phenyl-5-methyl-2-hexenal, 3-methyl-butanal, 2-methyl-butanal and 2-phenyl-2-butenal) (**Figure 4D**). Additionally, five compounds were uniquely detected in the AO_RIB40 fermentations, that did not pass the p-value cutoff < 0.01 but had p-values < 0.05 (heptanal, 4-methyl-3-pentenal, heptanol, ethyl phenylacetate and ethyl 2-methylbutyrate). Some compounds of note include 2,3-butanediol and R-2,3-butanediol, which were uniquely detected in AO_RIB40 and contributes to the floral and fruity aroma of soy sauce, 3-methyl-Butanal, which is an aldehyde detected in moromi fermentation [30], and 2-methyl-butanal, which has a cocoa or coffee-like aroma and has been detected in fermented sausages [31].

## Discussion

Food fermentation has a profound impact on the taste and flavor profile of foods. However, little work has compared the impact of *A. oryzae* and *A. flavus* on these compounds during food fermentation. To address this gap, we conducted rice fermentations using three distinct starter cultures: the food-grade strain AO_RIB40 and two closely related wild strains, AF_NPK13tox and AF_AflaGuard. We performed volatile compound profiling using DH-GC-MS and evaluated sensory attributes using an e-tongue (**Figure 1**). In addition, we quantified aflatoxin production to assess food safety and measured alpha-amylase activity to evaluate fermentation efficiency across the starter cultures.

The higher alpha-amylase output of AO_RIB40 (**Figure 2**) is a direct consequence of domestication, where artificial selection favored more efficient starch metabolism [4, 5, 10]. Consistent with a previous proteomic analysis of rice fermentation, AO_RIB40 exhibited significantly higher alpha-amylase activity than the *A. flavus* strains [4]. This parallels genomic observations that all *A. oryzae* isolates carry multiple copies of the α-amylase gene, whereas wild *A. flavus* typically has only one copy [4, 5]. The *A. flavus* strains in our study (with one alpha-amylase gene copy) showed significantly weaker amylase activity. The enhanced starch breakdown by AO_RIB40 likely accelerates saccharification and yeast fermentation, increasing ethanol yield and fermentation speed. Interestingly, variation in alpha-amylase activity is present even within *A. oryzae* strains. For instance, Chacón-Vargas *et al.* 2020 found that AO_RIB40 produced significantly more amylase (and grew faster on starch) than *A. oryzae* 14160 during solid rice fermentation [9], and suggested the importance of strain selection depending on fermentation substrate.

Additionally, we quantified aflatoxin production across our three isolates and confirmed that only AF_NPK13tox produced aflatoxin. AO_RIB40 has at least 4 loss-of-function mutations in the aflatoxin encoding gene cluster [32] and AF_AflaGuard has a large deletion encompassing the entire aflatoxin encoding gene cluster [25, 33]. However, despite this lack of aflatoxin production in AF_AflaGuard, this strain lacks the alpha-amylase activity and favorable sensory profile of AO_RIB40 (**Figures 3** and **4**).

The E-tongue results reveal clear differences in compounds associated with flavor attributes. Based on these results, it is anticipated that rice fermentations of *A. oryzae* RIB40 would be perceived has producing significantly less bitterness, astringency, and aftertaste A compared to those made with *A. flavus* strains (**Figure 3**). This observation likely reflects the different metabolite profiles between starter cultures and may be explained by a shift from primary metabolism in *A. oryzae* (e.g., high alpha-amylase activity) to secondary metabolism in wild *A. flavus* isolates. Secondary metabolites could lead to bitter compound production, and, generally, less proteolytic activity by shifting energy investment away from primary metabolism [34]. Due to the safety concerns with consumption, these E-tongue results provide the first insights into the potential flavor differences produced by these two strains.

Our study demonstrates that rice fermented with *A. oryzae* produces a significantly richer volatile profile in compounds associated with favorable sensory attributes compared to fermentations performed with *A. flavus* strains [35]. Specifically, *A. oryzae* fermentations yielded higher levels of unsaturated alcohols, ketones, and aldehydes that contribute complex sensory attributes [36]. Among the elevated compounds were 2-methyl-3-buten-2-ol and 3-octen-2-one. 2-methyl-3-buten-2-ol may impart distinctive fruity or green nuances that heighten the freshness of the aroma, while 3-octen-2-one is generally associated with mushroom or earthy notes, which may suggest a combination of earthy and umami attributes in the fermented rice [37].

Furthermore, both 2-methyl-butanal and 3-methyl-butanal (aldehydes associated with malty aromatic notes) were produced at significantly higher levels in the AO_RIB40 fermentations (**Figure 4D**). These compounds were also detected in rice koji fermented by *A. oryzae* [38]. These branched-chain aldehydes are key contributors to the characteristic rich malty aroma of soy sauce [39, 40].

Several heterocyclic compounds (4-methyl-1,3-dioxane and 2,4,5-trimethyl-1,3-dioxolane), were also more abundant in the *A. oryzae* fermentation. These compounds, often formed through Maillard-type reactions during fermentation, can contribute sweet, caramel-like, or even complex fruity notes that enhance the overall flavor profile [35]. Their presence may also indicate *A. oryzae*’s more efficient enzymatic transformations and sugar degradation processes [11]. We also detected several alcohols at significantly greater abundance in AO_RIB40 compared to the *A. flavus* fermentations (*i.e.* 2-methyl-3-Buten-2-ol, hexanol, 2-(2-Naphthyl)-2-propanol, 2-hexanol, pentanol, 2,3-butanediol, R-2,3-butanediol, and propanol). These observations are consistent with rice wine fermentation, where alcohols are more abundant early in the fermentation process and are gradually converted into esters and other sensory compounds during as fermentation continues [41, 42].

Additionally, phenyl-substituted aldehydes such as 4-methyl-2-phenyl-2-pentenal, 2-phenyl-5-methyl-2-hexenal, 3-phenyl-2-pentene, and 2-phenyl-2-butenal, as well as the aromatic alcohol 2-(2-naphthyl)-2-propanol, were present in higher concentrations. These compounds often contribute floral, balsamic, and nutty nuances that have been linked to elevated sensory appeal in rice-based fermentations, consistent with the role of aromatic ring structures in enhancing sweet and floral odor profiles [35]. The appearance of these compounds suggests that *A. oryzae* fermentations favor the release or synthesis of aromatic precursors derived from amino acid metabolism and lipid oxidation [43]. Further, higher abundances of simpler alcohols (2-hexanol, pentanol, and propanol), as well as branched aldehydes (3-methyl-butanal, 2-methyl-butanal, and isobutanal), were detected at higher abundance in the AO_RIB40 fermentations. These compounds are known to contribute green, fresh, malty, and even slightly fatty aromas, all of which are desirable in the context of rice fermentations for achieving a well-rounded flavor profile [35].

Our results are similar to Bodinaku *et al.* (2019), in which *Penicillium* strains were experimentally evolved in conditions mimicking the cheese making environment and volatile compounds were compared against the ancestral strains [44]. The wild ancestors characteristically produced musty, earthy odorants such as geosmin, whereas lineages serially cultured in the cheese environment largely lost the capacity to synthesize this compound.

Instead, cheese lineages shifted their metabolome and produced volatile compounds contributing descriptors like “cheesy” and “fatty,” which are desirable attributes in cheese making. This study, and our previous work [4], identified a down-regulation of genes involved in the production of secondary metabolites in strains adapted to the food environment compared to their wild counterparts, which likely indicates a shift away from the production of costly defense compounds not needed in the nutrient-rich, low-competition fermentation environments, where the energetic cost of producing a broad array of secondary metabolites is disadvantageous ([34, 45]).

### Conclusions

This study highlights the biochemical and sensory distinctions between the domesticated strain AO_RIB40 and its wild relatives, AF_NPK13tox and AF_AflaGuard, during rice fermentation. The enhanced α-amylase activity and absence of aflatoxin production in *A. oryzae* underscore its safety and effectiveness as a fermentation starter. Notably, rice fermented with *A. oryzae* exhibited a more favorable sensory profile, characterized by reduced bitterness and astringency and an increased umami-rich aftertaste. Volatile compound analysis revealed that *A. oryzae* produced significantly higher levels of alcohols, aldehydes, ketones, and heterocyclic compounds associated with fruity, malty, and umami aromas. Collectively, these findings suggest that long-term domestication has shaped the volatilome of *A. oryzae* to enhance desirable sensory qualities in fermented foods.

## Data availability

Raw whole-genome Illumina data for A. flavus NPK13tox is available through the NCBI Sequence Read Archive through BioProject accession number PRJNA1277813.

